# The Rapid Mechanically Activated (RMA) channel transduces increases in plasma membrane tension into transient calcium influx

**DOI:** 10.1101/2025.08.06.668926

**Authors:** Yannick Guerringue, Sebastien Thomine, Jean-Marc Allain, Jean-Marie Frachisse

## Abstract

- Plants respond to mechanical stimuli by a rapid increase in cytosolic calcium. The intensity and kinetics of the calcium changes define calcium signatures important for biological responses . In this study, we determine the properties of a calcium permeable force-gated channel localized at the plasma membrane called Rapid Mechanically Activated (RMA).
- Using patch-clamp and pressure-clamp, we characterized the kinetics of activation and inactivation of RMA channel upon stimulation by pulses of pressure applied onto the plasma membrane. Combining repetitive pressure pulse protocols at different frequencies with modeling, we investigated the channel’s capacity to transduce high frequency mechanical stimuli.
- RMA channel rapidly activates in response to membrane tension, then it inactivates during prolonged stimulation. Upon repeated stimulations, RMA current amplitude decreases irreversibly indicating that undergoes adaptation. The channel kinetics may be modeled with four chemical states and the model predicts that it behaves as a pass band filter in the 10 Hz - 1 kHz range.
- In conclusion, due to its activation/inactivation characteristics, RMA channel is a candidate to mediate cytosolic calcium signaling in response to mechano-stimulation. Its adaptation and filtering properties suggest its involvement in the transduction of high frequency mechanical stimulation such as those produced by insects’ vibrations.

## Introduction

Plants respond rapidly to mechanical stimulation by an increase in cytosolic calcium concentration activating signalling pathways via calcium-binding proteins (Monshausen *et al*., 2009) (Dodd, Kudla and Sanders, 2010). A high diversity of calcium signals has been monitored, depending on the type of stimulation or the selected tissue (McAinsh and Pittman, 2009; Martí, Stancombe and Webb, 2013). The spatial and temporal dynamics of these calcium signals are specific of the stimulus that initiated them, so that they were called calcium signatures. Although the recording of calcium signals and the identification of calcium transporters have been carried out for more than two decades, the channels responsible for the transduction of mechanical stimulations into calcium entry in the cytosol still need to be identified (Wilkins *et al*., 2016). In this context, it is necessary to further characterize the gating properties of calcium channels, their mode of activation, their kinetics and their conductance to understand their contribution to calcium signatures.

Mechanosensitive channels represent ideal candidates for mechanotransduction as they are activated by mechanical stress and then directly transduce mechanical stimuli into ion fluxes (Peyronnet *et al*., 2014; Guichard, Thomine and Frachisse, 2022). The detailed investigation of the gating mechanism of mechanosensitive channels such as the bacterial MscL and MscS or the mammalian TREK/TRAAK K_2P_ and Piezo allowed understanding how their activity is tuned by the mechanical tension of the membrane (Douguet and Honoré, 2019; Martinac 2011). Compared to these channels, the description of the gating mechanism of plant mechanosensitive channels is still in its infancy. In plants, genes encoding mechanosensitive channels have been identified by homology with bacterial and animal genes such as the MSL family or Piezo (Haswell, 2007; Zhang *et al*., 2019). MCA and OSCA genes encoding mechanosensitive channels have been discovered on the basis of the altered calcium signalling observed in the corresponding mutants (Nakagawa *et al*., 2007; Yuan *et al*., 2014). Their mechanosensitivity has been verified for three MSLs (Mechanosensitive Small conductance-Like) and five OSCAs expressed in oocyte and human HEK cells respectively (Maksaev and Haswell, 2012; Hamilton *et al*., 2015; Lee *et al*., 2016; Murthy *et al*., 2018). Moreover, MCA2, when reconstituted in lipid bilayers, also showed a force gated activity (Yoshimura, Iida and Iida, 2021). However, very few endogenous mechanosensitive channels have been studied in plant cells, even though their accurate characterization should be performed in their native membrane.

A mechanosensitive channel localized at the plasma membrane of Arabidopsis cells was recently discovered by Tran et al. (2017) and called Rapid Mechanically Activated (RMA). It was reported as a Ca^2+^-permeable mechanosensitive channel, characterized by rapid activation and inactivation. RMA currents have been found to be dependent on the transmembrane protein DEK1. DEK1 is predicted to be composed of a 23 transmembrane helix-domain and a cytosolic C-terminal domain, which is homologous to animal protease calpain (Kumar, Venkateswaran and Kundu, 2013). The deletion of the CALPAIN domain is embryo-lethal while the deletion of the transmembrane domain disturbs the mechanosensitivity of the RMA channel (Tran *et al*., 2017a). It was not possible for the authors to specify whether or not DEK1 constitutes a subunit of the channel and they proposed that DEK1 is a regulator of RMA that modifies its gating properties (Guerringue, Thomine and Frachisse, 2018; Malivert, Hamant and Ingram, 2018).

RMA represents the first plant Ca^2+^-permeable mechanosensitive channel that has been recorded in its native membrane. In order to improve the knowledge about this channel, we have performed a thorough characterization of its activity. We analysed its gating properties in relation with mechanical tension applied to the membrane and its activity-dependent adaptation. In our study, we have developed a model of activation supported by a computational analysis, which highlights the peculiar features of the channel. We propose, in this model, two possible modes of channel adaptation. One that is pressure-dependent arising from the membrane-channel interaction, and another that is pressure-independent implying a biochemical reaction linked to a particular state of the channel. We also investigated how the channel responds to repetitive stimuli such as found in nature, like acoustic signals from bees or vibrations from insect chewing and generalist herbivore (Appel and Cocroft, 2014; Pinto *et al*., 2019; Veits *et al*., 2019). The model we developed suggests that the channel may act as a detector of oscillations in a specific frequency range. Finally, our study provides a description of the RMA Ca^2+^-permeable channel that helps to understand how calcium signals are generated during mechanotransduction, and provides elements to further identify the channel and study its regulation.

## Materials and methods

### Callus generation

Surface-sterilized seeds from Arabidopsis (Arabidopsis thaliana) Columbia-0 (Col-0) wild-type were sown on “initiation medium” containing 4.3g/L Murashige & Skoog salts (MS, Sigma-Aldrich), 2% sucrose, 10mg/L Myo-Inositol, 100µg/L Nicotinic Acid, 1mg/L Thiamine-HCl, 100µg/L Pyridoxine-HCl, 400µg/L Glycine, 0.23µM Kinetin, 4.5µM 2,4-D, 1% Phytagel, (pH 5.7). For callus generation, seeds were cultured in a growth chamber for 15 days. Calli were then transferred onto “maintenance medium” containing 4.3g/L MS salts (Sigma-Aldrich), 2% sucrose, 10mg/L Myo-Inositol, 100µg/L Nicotinic Acid, 1mg/L Thiamine-HCl, 100µg/L Pyridoxine-HCl, 400µg/L Glycine, 0.46µM Kinetin, 2.25µM 2,4-D, 1% Phytagel, (pH 5.7), and sub-cultured every 15 days onto fresh “maintenance medium”.

### Protoplast preparation

Calli were digested for 15 min at 22°C under hyperosmotic conditions (2mM CaCl2, 2mM MgCl2, 1mM KCl, 10mM MESs (pH 5.5), 0.2% Cellulysin (Calbochem®), 0.2% cellulase RS (Onozuka RS, Yakult Honsha Co.), 0.004% pectolyase Y23 (Kikkoman Corporation), 0.35% Bovine serum albumine and mannitol to 600 mOsmol. For enzyme removal, the preparation was washed twice with 2mM CaCl2, 2mM MgCl2, 10mM MES (pH 5.5), and mannitol to 600 mOsmol. For protoplast release, the preparation was incubated with 2mM CaCl2, 2mM MgCl2, 10mM MES (pH 5.5), and mannitol to 280 mOsmol. The suspension was filtered through a 50 µm nylon mesh (Tran *et al*., 2017a).

### Patch clamp

Patch-clamp experiments were performed at room temperature with a patch-clamp amplifier (model 200A, Axon Instruments, Foster City, CA) and a Digidata 1322A interface (Axon Instruments). Currents were filtered at 5 kHz, digitized at 20 kHz, and analyzed with pCLAMP8.1 and Clampfit 10 software. During patch-clamp recordings, cells were held at a holding potential (not corrected from liquid junction potential of -5 mV for pipette/bath solutions Cs2SO4 75 mM/CaCl2 50 mM) of − 180 mV and pressure was applied with a High Speed Pressure-Clamp system (ALA Scientific Instrument, NY), allowing the application of precise and controlled pulses in the pipette. For Ca2+ current recordings, bath solutions contained (mM): 50 CaCl2, 5 MgCl2, 0.25 LaCl3 and 10 MES-Tris (pH 5.6); while pipettes were filled with (mM): 75 Cs2SO4, 2 MgCl2, 5 EGTA, 4.2 CaCl2, and 10 Tris-HEPES (pH 7.2), supplemented with 5 MgATP. To exchange Ca2+ against Na+, a bath solution containing (mM): 100 NaCl, 5 MgCl2, 0.25 LaCl3 and 10 MES-Tris (pH 5.6) was used.

Osmolarity was adjusted with mannitol to 450 mOsmol for the bath solution and to 460 mOsmol for the pipette solution using an osmometer (Type 15, Löser Meβtechnik). Gigaohm resistance seals between pipettes (pipette resistance, 0.8-1.5 MΩ) (coated with Sylgard (General Electric) pulled from capillaries (Kimax-51, Kimble Glass)) and protoplast membranes were obtained with gentle suction leading to the whole-cell configuration, then excised to an outside-out configuration. The current inactivation kinetics were fitted with a mono-exponential function: F(t)= A * e-t/τ + C, where A is the coefficient, τ is the time constant and C represents the maximum current intensity.

### Modelling

See Supplement data S4: Equations and constants of the model.

## Results

### Dynamic properties of RMA during a square pulse

The characterization of the RMA conductance has been carried out in the experimental conditions previously described by (Tran *et al*., 2017a). Outside-out excised patches from protoplasts extracted from *Arabidopsis thaliana* calli were subjected to pressure pulses up to 200 mmHg (Fig.1A). The extracellular bath solution contained Ca^2+^ as the major cation whereas, unlike in Tran et al. (2017), the main anion in the pipette solution was SO_4_ ^2-^, which has the advantage to be non-permeant for the MSL mechanosensitive channels and then to exclude their activity from our recordings.

**Figure 1.**
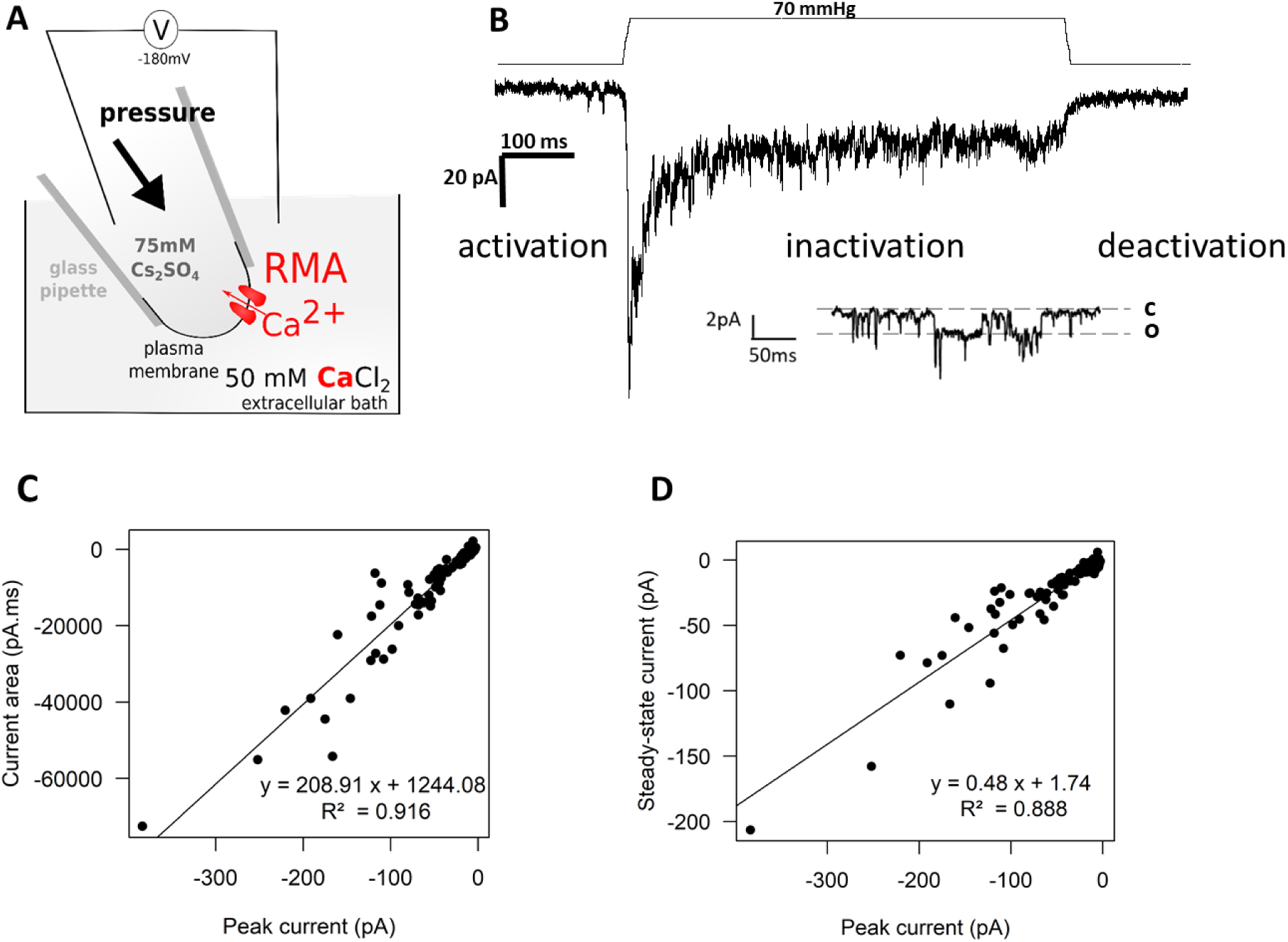
Kinetics of the RMA conductance under a square pressure pulse. (A) Experimental conditions for recording the RMA conductance. The plasma membrane of a protoplast was sealed to a glass pipette (GigaOhm seal) and a small patch of membrane was excised in the outside-out configuration. Pipette solution contained mainly 75 mM Cs_2_SO_4_, bath solution was mainly composed of 50 mM CaCl_2_, for detail see Materials and Methods. (B) Recording of the RMA conductance under a square 70 mmHg-pressure pulse. A single-channel recording taken from another trace shows the closed state *c* and the open state *o* of the channel. (C-D) Current area (C) or steady-state current (D) versus peak current (n = 10). Linear correlation fit was represented by a straight line. The equation and the correlation coefficient are shown on the graph.

Excised patches in the outside-out configuration were clamped at -180 mV and submitted to square pulses of increasing pressure, which triggered a negative current (Fig.1B) that was previously shown to correspond to an entry of Ca^2+^ (Tran *et al*., 2017a). Indeed RMA was reported to permeate the divalent cations Ca^2+^ and Ba^2+^. To check whether this channel is also permeant to monovalent cations, the RMA conductance was recorded first in standard conditions with 50 mM Ca^2+^ in the bath, then Ca^2+^ was substituted with 100 mM Na^+^. The substitution of the Ca^2+^ by Na^+^ (same equivalent charge) in the bath solution did not alter the intensity of the peak current, whatever the pressure or the voltage (Fig. S1). Therefore, the conductances of RMA to Ca^2+^ and Na^+^ seem close. This result indicates that RMA is a cationic non-selective channel like Piezos (Coste *et al*., 2012) and OSCAs (Murthy *et al*., 2018).

Pressure pulses triggered a fast current increase, corresponding to the activation of RMA channels, rapidly followed by a decrease, while pressure was maintained (Fig.1B). A steady state current remained which returned to the initial value after the end of the pressure pulse. These dynamics suggested a basic three-state model with closed, opened and inactivated channels. The transition from open to closed channels at the end of the pressure pulse was called deactivation while the transition from open to inactivated channels upon sustained pressure was called inactivation. The peak current, the area under the curve and the steady-state current were measured at different pressure values and we found a linear correlation between the peak current and the area under the curve or the steady-state current (Fig.1C-D). The relationship between the peak current and the steady-state current was pressure-independent, suggesting that the inactivation is pressure-independent. Therefore, we chose to use the peak current as a parameter of activation of RMA.

The rates of activation and deactivation were in the same range or faster than the response of the pressure-clamp system, in the order of a few milliseconds (Fig.S2). Hence we could not determine the pressure dependence of these parameters. Nevertheless, as the activity of RMA is increased by an increase of pressure, the activation-deactivation kinetics must be pressure-dependent. We therefore propose a gating model for RMA composed of 3 states (closed, opened, inactivated) where the activation-deactivation kinetics are pressure-dependent.

### RMA activity upon repeated stimulations

RMA activity decreases upon repetitive activation. This decrease, often called rundown, is frequently encountered in many patch-clamp experiments even though it is seldom mentioned in the literature. We studied in detail this phenomenon, that we called adaptation, so as to include it in our model, our perspective being to understand the cause of the adaptation. To do so, we applied the same pulse of pressure to the patch of membrane and measured the peak current (Fig. 2A). The time interval between two stimulations was changed from 1 to 120 s to discriminate between the effect of time and the effect of the number of stimulations. The decrease of activity was much faster when the pressure pulses were every 1 s compared to every 120 s (Fig.2B). In contrast, there was no difference in the rate of decrease of activity at different time intervals (1s, 30s or 120s), when it was expressed as a function of the number of pulses (Fig.2C). These results indicate that the decrease of activity is dependent on the number of stimulations rather than on time. The decrease in the intensity of the peak current followed an exponential decay that we fitted with an exponential function for each patch so as to quantify its kinetics. The time constant of the fit, which we called the adaptation constant, was correlated to the time interval between stimulations (Fig.2D), confirming that the decrease of activity was not time-dependent but stimulation-dependent. Adaptation was irreversible in the time frame of our experiments, as a gap of 30 to 45 min with no stimulation was no sufficient to restore initial activity. Surprisingly, the adaptation constant was not dependent on the duration of the pressure pulse (Fig.2E-F) suggesting that it happens in the very first instant of the pulse.

**Figure 2.**
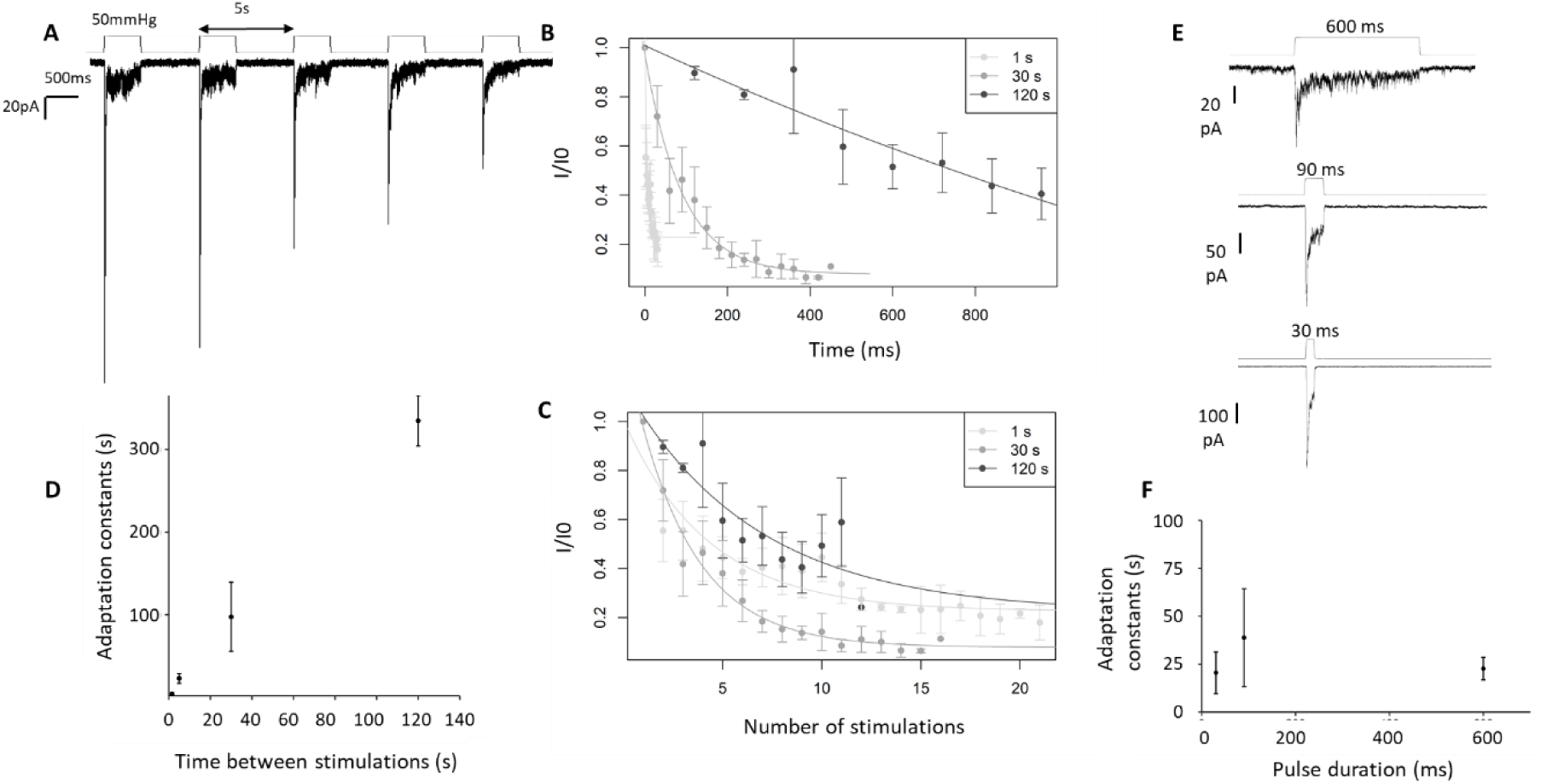
The activity of RMA decreases upon repetitive stimulations. (A) Recording of the RMA conductance under repetition of square pulses at 50 mmHg every 5 s. (B-C) The same protocol has been carried out with different time intervals between stimulations. Mean (± SE) of the peak current *I* normalised by the first peak current *I_0_* over time (B) and over stimulations (C) for time interval of 1, 30 or 120 s and a duration of stimulation of 600 ms (n ≥ 5). (D) The individual decay curves have been fitted with an exponential function from which we deduced the time constant (adaptation constant). This adaptation constant is presented with different time intervals (n ≥ 5). (E) Recordings of the RMA current under a square pulse of pressure with different duration of stimulation of 600, 90 or 30 ms. (F) Adaptation constant for different pulse duration (n ≥ 3).

Together, these results show that RMA adaptation is driven by the number of mechanical stimulations but independent of the time or the duration of the stimulations in the range used in this study.

### RMA is activated at a high threshold of membrane tension

To determine the activation curve of RMA, square pulses of increasing pressure were applied on outside-out patches of membrane until the breaking of the patch. The sensitivity to pressure of RMA current was variable and dependent of each patch as exemplified in figure 3A with four recordings obtained from different patches activated by the same pulse protocol. The peak current was measured for each pressure and the activation curve was plotted and fitted with the following Boltzmann equation (Fig. 3B):

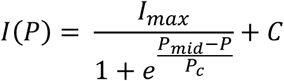

**Figure 3.**
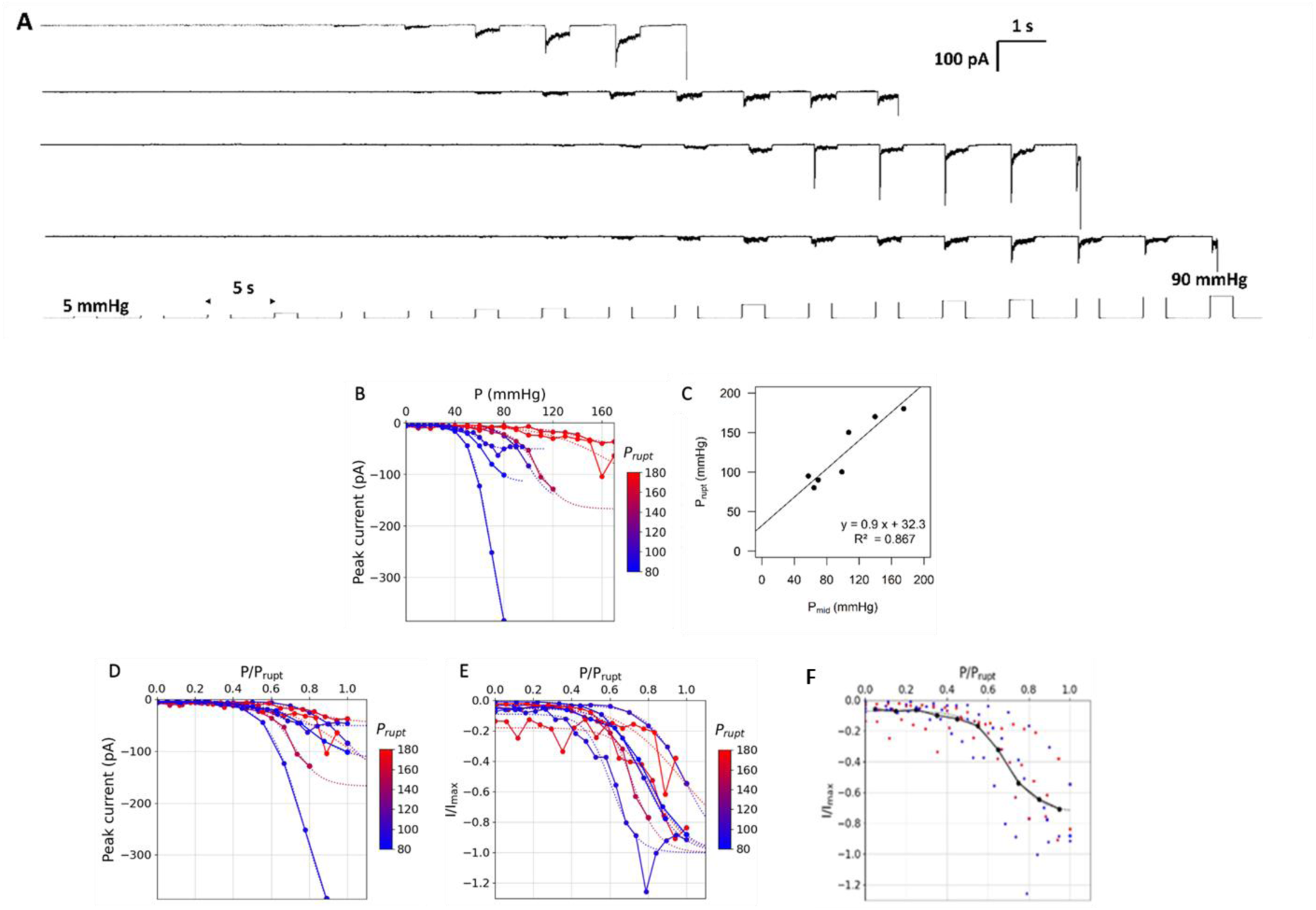
RMA activation threshold correlates with the membrane rupture pressure. (A) Recordings of the RMA conductance in four independent patches under square pulses of increasing pressure until the patch rupture. (B) Peak current at different pressure values. Each curve corresponds to one independent patch is colored according to the rupture pressure *P_rupt_* (in mmHg) and is fitted (dotted line) by a Boltzmann equation. (C) The rupture pressure *P_rupt_* was correlated to the pressure at mid-activation *P_mid_*. Linear correlation fit was represented by a straight line. The equation and the correlation coefficient are shown on the graph. (D) Peak current in function of *Pressure*/*P_rupt_*, each curve is fitted by a Boltzmann function. (E) Normalized peak current in function of *Pressure*/*P_rupt_*, each curve is fitted by a Boltzmann fnction. (F) Fit of the mean curve with a Boltzmann function which P/Prupt at mid-activation is 0.73 .Each curve (n = 7) corresponds to one independent patch, the peak current was normalised by the maximum current I_max_ and the pressure was normalized the rupture pressure P_rupt_.

*I_max_*, representing the total current when all the channels in the patch are open, had an average value of -154 pA ± 138 pA (n=7), corresponding to the opening of around 75 channels (Tran *et al*., 2017a). *P_mid_*, the parameter of sensitivity to pressure, had an average value of 97 mmHg ± 32mmHg, which is high compared to the anion permeant mechanosensitive channel MSL10 *P_mid_* of 49.3 ± 3.4 mmHg co-residing on the same membrane and characterized in the same conditions (Tran et al., 2021). The high variability in the parameters suggested that P is not the parameter gating RMA. We noticed that the higher the pressure to activate RMA channels was, the higher P_mid_ was and the higher the pressure P_rupt_ to break the patch was (Fig. 3A and B). P_mid_ was correlated to P_rupt_, the pressure at which the patch ruptured (Fig.3 C). Moreover, we also observed that P_first peak_, the lowest pressure triggering inactivating RMA channel activity, is correlated with P_rupt_ and P_mid_ (Fig. S3 A and C). If we assume that the rupture of the patch occurs at a constant tension, this suggests that the activation of RMA depends on membrane tension rather than on pressure. Tension is related through Laplace’s law to pressure via the radius of curvature of the patch. Depending on the geometry of the patch, the same pressure would not trigger the same membrane tension, which accounts for the high variability in pressure sensitivity (Lewis and Grandl, 2015; Bavi *et al*., 2016). When RMA activity was ploted as a function of *P*/*P_rupt_*, *P*/*P_rupt_* = 1 corresponding to the patch rupture, as an estimate of membrane tension, the variation among the activation thresholds obtained from different patches decreased significantely (Fig. 3D). This further support that RMA is responsive to tension rather than pressure. As the rupture pressure P_rupt_, the pressure at first peak P _first peak_ could be used for the estimation of the membrane tension, with the advantage that determining P _first peak_ in not destructive (Fig. S3).

Moreover, since the total activation depends on the number of channels in the patch, the peak current was normalised by the maximum current *I_max_* (Fig. 3E). The average ratio *I*/*I_max_* was then plotted over the ratio *P*/*P_rupt_*, which is considered as an estimate of membrane tension (Fig. 3F). The mean curve was fitted with a Boltzmann function which *P*/*P_rupt_* at mid-activation was 0.73. This value, close to the membrane rupture, means that the sensitivity of RMA to membrane tension is quite low. Therefore, RMA channels are activated at membrane tensions close to the rupture.

### Characterization of RMA inactivation

During a pressure pulse, the rapid activation of RMA channels was quickly counterbalanced by their inactivation, which shapes the signal as a peak. Although the peak current was dependent on tension, it did not seem to be the case for RMA inactivation. Indeed, the time constant of current decay, determined by a monoexponential fit (Fig.S2B), was constant over *P*/*P_peak_* (Fig.4A). The time constant ranged between 10 and 50 ms and was tension insensitive. Moreover, measurement of the inactivation time constant at different voltages indicated that RMA inactivation kinetics is not affected by membrane potential (Fig.4B). Hence inactivation of the RMA channels is tension- and voltage-independent.

**Figure 4.**
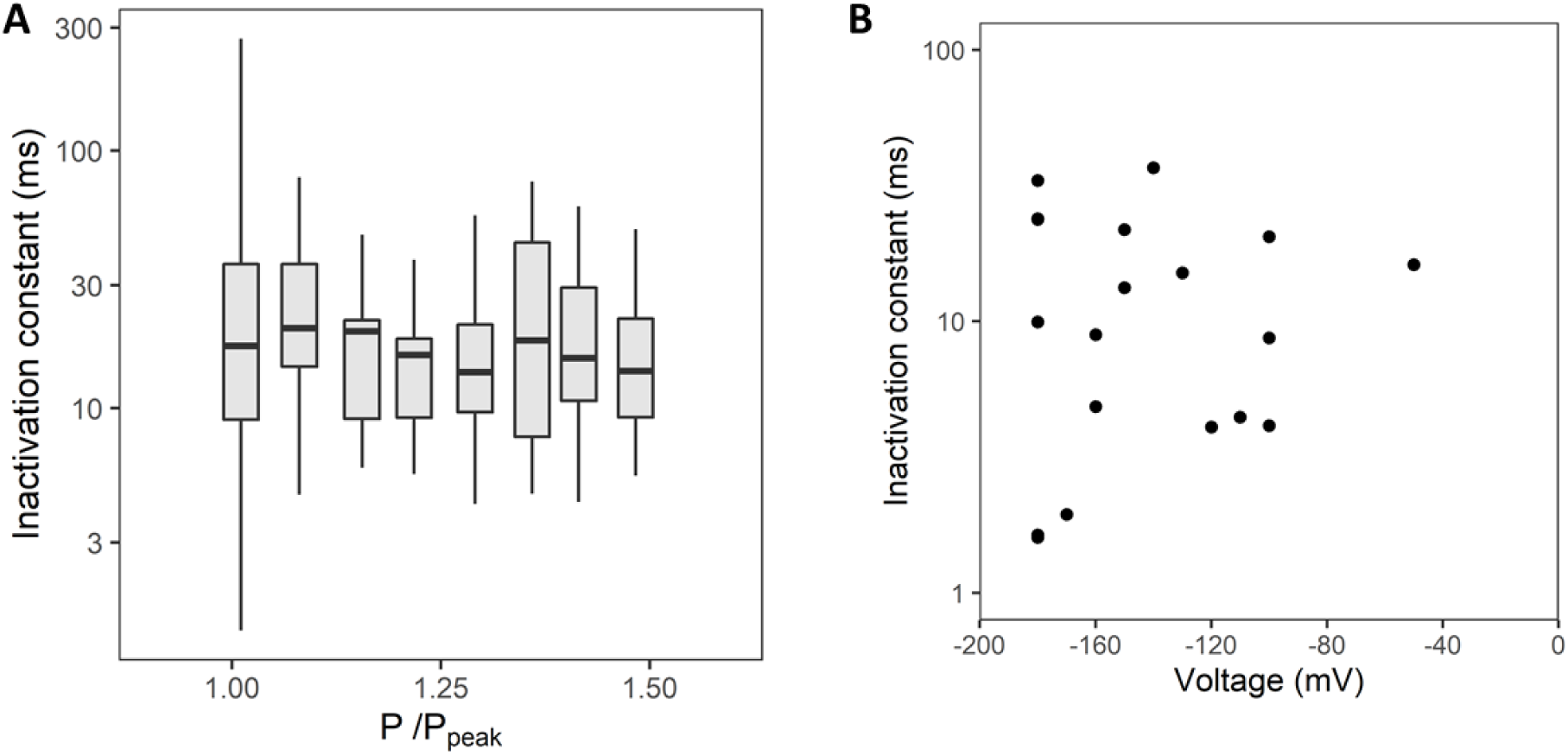
Characterization of the RMA inactivation. (A) The current decrease under a pressure pulse was fitted with an exponential function. The time constant of this fit, named inactivation constant, is presented for different values of *P*/*P_peak_* used a proxy for the membrane tension in the patch (n = 28). (B) Inactivation constant over voltage (n = 3).

### Construction of the four-state gating model of RMA

The model we propose for RMA aims to capture all the described effects while minimizing the number of parameters. Our model is based on the assumption that RMA channel has well-defined chemical states (closed, open, inactivated and adapted). The changes of channel configuration are defined by transition rates. Mechanosensitivity can then be modeled as a pressure-dependency of the transition rate following a Bell’s law (Tran *et al*., 2021).

A minimal model for channel gating only includes a closed state (denoted C) and an open state (denoted O). However, such a model cannot capture the inactivation of the channel. When an increment of pressure is applied, the open state is favored, leading to a variation of the current. This current will gradually increase to a plateau value and remains stable (see for ex. (Tran *et al*., 2021) for such case). Therefore, an additional inactivated state (denoted I) is needed.

We tested three possible hypotheses for the transition to the inactivated state (see Supplementary Figure S4). The simplest one was that the inactivation process corresponds to a transition from the open state only (meaning channels can move from O to I - and back to O). This reflected well the inactivation of the channel, but not its slow adaptation. The second hypothesis corresponded to a transition from the close state to the inactivated state only (meaning from C to I, and back to C), with the underlying hypothesis that the adaptation may be due to an inactivation of the closed channels. This alternative however does not allows recapitulating both the inactivation and the adaptation. A third alternative is the classical four-state model of Hodgkin-Huxley (Hodgkin and Huxley, 1952), with an inactivated state associated with the closed state Ic, and one with the open state Io (and a transition between the two). However, the choice of transition rates based on their constraints imposed by the thermodynamical balance leads to a model where the Ic state is almost never significantly filled. Hence, this model behaves very much as the simplest 2-state model. Finally, the first model that associates inactivation with the open state best recapitulated the experimental results, although it does not account for adaptation (Fig.5A). In agreement with the experimental data, we did not introduce a pressure-dependency of the inactivation transition rates. Taken together, this implies that inactivation is not related to the membrane tension, but rather to an inner process of the channel.

**Figure 5.**
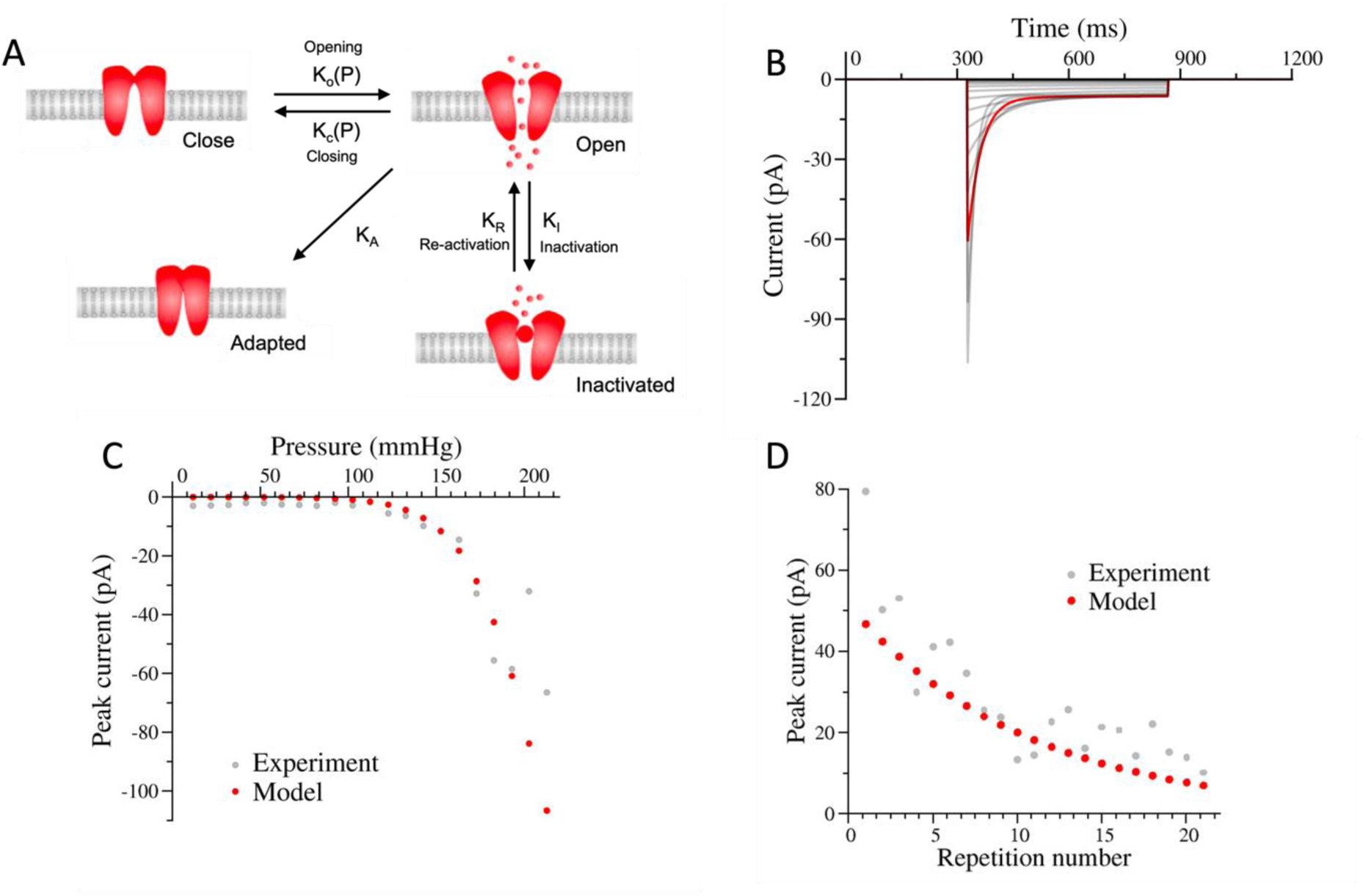
Model of activation of the RMA channel. (A) The four-state gating model of RMA: close, open, inactivated and adapted. (B-D) simulation of the current when a patch is exposed either to a protocol of pulses of increasing pressure (B,C) or a protocol of repeated pulses at constant pressure (D). Simulated data in red are overlayed on experimental data in grey. (B) current obtained with increasing pulses of pressure. (C) current at the peak as a function of pressure. (D) current at the peak as a function of the number of pulse stimulation of 183mmHg

Since the model could not capture the adaptation (see Fig. 2), we added a fourth “adapted” state (denoted A, Supplementary Data S5). Channels in this state remain closed and do not return to other states within the time range of our experiments, thus it is modeled as an irreversible transition. Stimulation dependency can be obtained by introducing a tension-dependent acceleration of the transition rates to the adapted state from all other states. Another option is to keep the transition to the adapted state tension-independent, but to have different rates from each state. Although a tension-dependent transition to adapted state from all states fits the experimental data, the most parsimonious model to couple adaptation to stimulation is to have a transition only from the open state to the adapted state, with a constant rate (see Fig.5A). We present the simulations using this last model, keeping in mind that others choices were possible.

Figure 5 shows the results of the simulation for a typical RMA channel. Figure 5A summarizes our model with the different transitions. Figure 5B shows the current vs time for increasing pressure steps. As expected, it reflects well the inactivation: the current displays a very fast peak, followed by a decrease of the ion flow. The brevity of the initial peak is such that it is poorly resolved, and the opening rate can be determined only approximately. However, the inactivation behavior is well described, as well as the stationary current. We used here the Pressure to mimic the experimental condition, but the channels are more likely activated by the membrane tension (Fig. 3). We have assumed here that the pressure and the tension are proportional, through a constant which would have to be determined for each experiment (Lewis and Grandl, 2015; Bavi *et al*., 2016). As the model does not include noise, our plot is smoother than the experimental data.

Figure 5C shows the associated peak current vs pressure for increasing pressure steps, simulated and experimental. The shape is similar to a standard “Boltzmann” response, despite the unexpected decrease of the current for high pressure. This effect could be due to adaptation, although we may not exclude noise in the experimental records: as the peak of current is very narrow, it is sometime difficult to measure it precisely. Note also that, at very low pressures, the measured peak is not distinguishable of the signal noise, while the model can go to zero.

To characterize the adaptation shown in Figure 2, repeated pressure steps were simulated. Figure 5D shows the evolution of the peak intensity with repeated pressure steps, experimentally and computationally. The model simulates well the adaptation. Note that the computation includes the waiting time in which no pressure is applied: during that time, it is possible for inactivated channels to move back to open (and then close) states, and a tiny fraction to move to the “adapted” state. This implies that the parameters are the same in Fig. 5B, C and D.

The model we developed was able to simulate most of our experiments always using the same rate constants. However, we had to adjust the total number of channels (as it changes in each patch) and the pressures sensitivities, as the channel is not likely to be sensitive to pressure itself, but to the membrane tension, which is pipette and patch geometry-specific (Lewis and Grandl, 2015; Bavi *et al*., 2016). We obtained an inactivation rate around 10ms, in agreement with our experimental observation (see Figure 3A). The other constants are given in supplementary data (Supplementary Data S5), but are less relevant, as we assume instantaneous changes of pressure, while real changes may be in the order of magnitude as the open time constant.

### The RMA channel behaves like a band-pass filter

Plants are oscillating due to variations of their environments (wind, insect contact, etc.), and have to adapt to these changing environments (Appel and Cocroft, 2014; Veits *et al*., 2019; Tran *et al*., 2021). We used our model to predict RMA response to stimulation at different frequencies. Frequency response can be easily tested in the model using an oscillatory pressure *P* = *P*_0_ + Δ*P* sin 2*πft*, where *f*is the frequency, *P*_0_the average pressure, and Δ*P* is the amplitude of pressure oscillations. To determine whether the oscillations potentiate or depress the mean RMA signal, we compared the signal to the one without oscillations, at the static pressure *P*_0_, with the signal over an entire number of oscillations. To study the effect of oscillations in a range of pressure where the channel has high dynamic range, we set *P*_0_ close to *P_mid_*. We removed the Adpated state from the model – thereby reducing the number of states to three – as the signal decrease due to adaptation would make the comparison difficult.

Figure 6 shows the mean relative intensity versus the frequency, for different durations of the oscillations. We observe that the channel behaves as a band pass filter: the response is potentiated in a broad range of frequencies (around 10Hz to 10kHz). At higher frequencies, the oscillations induce a slightly higher mean signal than the static stimulation, while at lower frequency the oscillations are detrimental. This double effect arises from the 3-states nature of the channel, with fast close-open transition and slow inactivation. Indeed, 2-states channels exhibit only a low-pass behavior (Tran *et al*., 2021). Such a behaviour is recapitulated when the duration of the stimulation is much shorter than the inactivation time (1 ms). Note that, due to the progressive adaptation, the channel response at a given frequency would decrease in average when the duration of oscillation increases. This means that the channel may act as a detector of oscillations in a specific range of frequencies and durations.

**Figure 6.**
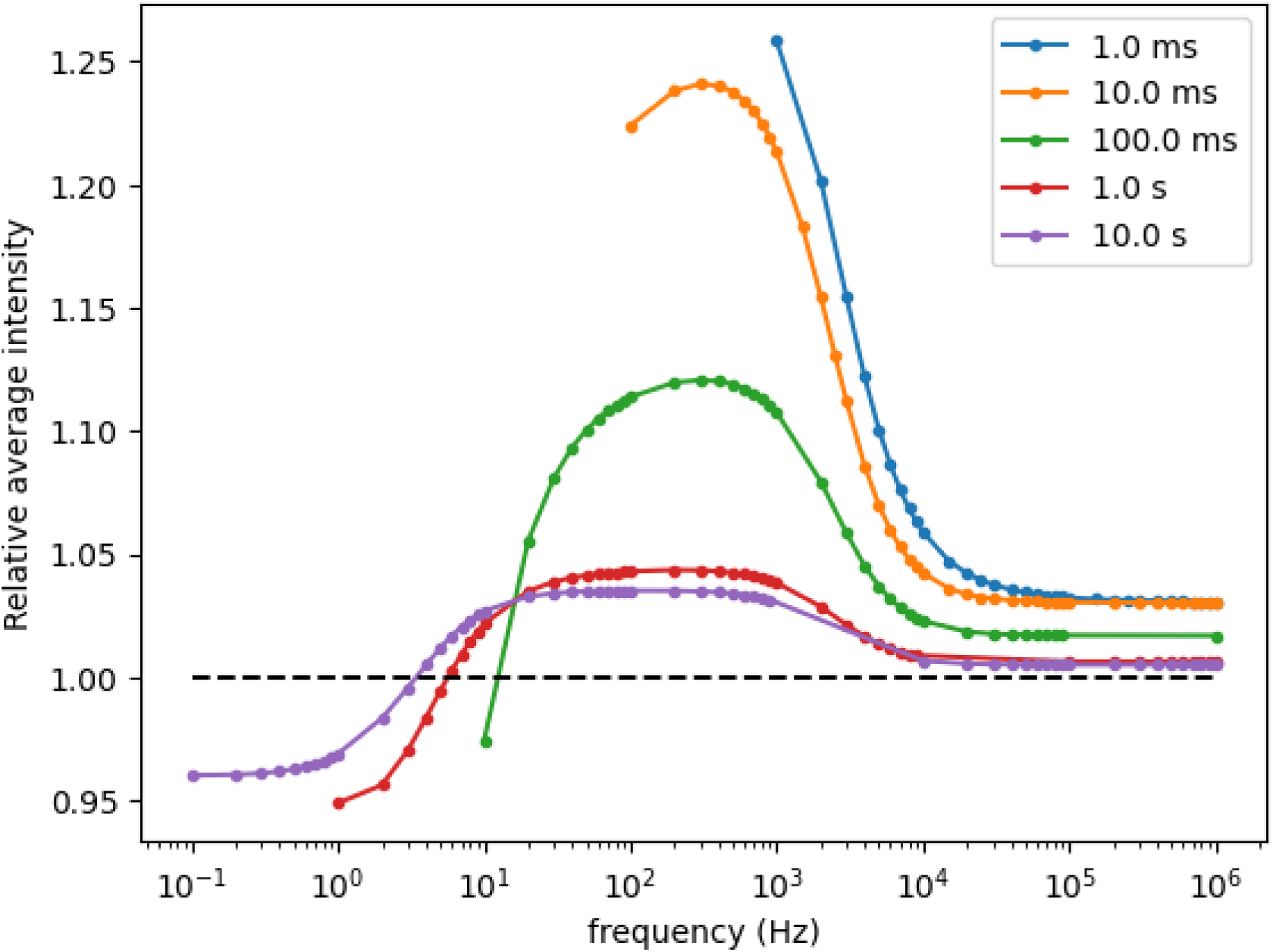
The RMA channel behaves like a band-pass filter. Average intensity for an oscillatory relative to a static pressure load as a function of frequency, for different durations of oscillations. The intensity is averaged over the duration of oscillations, and divided by the intensity at the static mean pressure. Parameters of the simulations are the ones from our experimental data (see Supplementary Table 1), with *P*_0_ = 175mmHg and Δ*P* = 25mmHg.

## Discussion

### RMA channels activate in response to membrane tension

Our thorough electrophysiological study allowed identification and quantification of the properties of the RMA current. Namely, we characterized its kinetics of activation, inactivation, deactivation, adaptation and the relationship pressure-activity. Under pressure clamp, the parameter controlled by the experimenter and applied on the membrane is the pressure. Analyzing the channel properties as a function of the pressure applied to the patch led to high variability (Fig 3), suggesting that tension rather than pressure might be the parameter controlling RMA activation. To directly assess membrane tension, it is necessary to accurately determine the curvature of the membrane at the pipette tip to apply the Laplace law (Lewis and Grandl, 2015; Bavi *et al*., 2016). However, measuring membrane curvature was not possible with our set-up. More generally, it is notoriously difficult to access to the tension of the membrane when applying the patch clamp technique (Sachs, 2015). It has been reported that the degree of stress of the membrane patch is likely to differ when comparing cell-attached patches to excised patches (Bavi *et al*., 2014, 2016). Indeed, the excision of the patch induces the rupture of the membrane continuity. Therefore, a significant stress profile difference (∼30%) is observed between inner and upper membrane leaflet in the excised configuration (Bavi et al., 2014). Neglecting a constant tension within the patch could lead to an underestimation of the pressure/tension efficient to activate MS channel. Therefore, to estimate membrane tension, we hypothesized that membrane rupture and the elicitation of the first peak occur at a constant tension. Normalization of the pressure either with the pressure inducing membrane rupture or with the pressure of the first peak appearance showed a consistent relationship with RMA activation across all membrane patches analyzed. The membrane rupture tension has been determined at 4 mN.m^−1^ on rye protoplast (Kell and Glaser, 1993). Based on this value, we could calculate a Tension_mid_ of ∼ 1.6 mN.m^−1^ and a Tension of ∼ 3.2 mN.m^−1^ for maximal RMA activation, which is close to rupture tension. Together, our results indicate that RMA, as other MS channels, is sensitive to membrane tension and not to pressure, with a high tension threshold.

### Adaptation

We have uncovered two significant properties of the RMA channel, (1) at the onset of a pressure pulse the channel activity increases rapidly. However during sustained pressure, channel activity decreases due to the transition to an inactivated state (Frachisse et al. 2020), (2) repeated stimulations result in a decrease in the number of activated channels. We have termed this process adaptation and propose that the transition to the adapted state occurs from the active state of the channel. To explore dependence of channel activity on membrane tension, we applied pressure up to high values on outside-out membrane patches. This configuration was chosen because it allowed the obtention of tight seals (GΩ), which was rarely the case in the cell-attached configuration. Patch excision may cause the loss or deterioration of cellular compounds involved in the regulation of mechanosensitive channel causing time-dependent run-down of channel activity after excision. While most MS channels are gated directly by the force from the surrounding lipids, some are tethered to subcellular structures such as proteins of the extracellular matrix or the cytoskeleton (Battle *et al*., 2015; Teng *et al*., 2015; Bavi *et al*., 2016). For RMA, the above hypothesis cannot account for adaptation because the decrease in activity would be time dependent rather than stimulation-dependent.

Adaptation of calcium signaling has been reported in more integrated biological systems. Allen et al investigated Ca signaling in guard cells that control the opening of stomata regulating CO_2_ and H_2_O fluxes at the interface between leaves and the atmosphere (Allen *et al*., 2001). Upon application of hypo-osmotic and hyperpolarizing conditions, the authors recorded an elevation in intracellular Ca^2+^ concentration, which they interpreted as being due to calcium permeable channel activity. When performing repetitive hypo-osmotic shock on guard cells, they noticed a decrease of the cytosolic calcium signal as stimulation progressed, which they called attenuation. Recently Audemar et al. (Audemar, Guerringue, Frederick, Vinet, *et al*., 2023) showed that repetitive mechanical stimulation of the Arabidopsis root also leads to rapid attenuation of calcium elevation. Changing the frequency and duration of stimulations, they concluded that the relevant factor for adaptation is the number of stimulations rather than their frequency. Then, RMA is a good candidate to mediate calcium signals that attenuate upon repetitive mechanical stimulations as those observed in guard cells and roots.

### The input of modeling

Using our modeling approach, we were able to determine the minimal components needed to capture the response of the channel. Four states are needed (see Figure 5A): one open state allowing ion passage through the membrane and 3 closed states with distinct properties. The deactivated closed state is reached when the tension activating the channel is released. The inactivated closed state is reached when the activating tension is maintained and the adapted closed state is also reached from the open state but this transition is irreversible. These states were deduced based on the experimental observations, the model enables to test the transition between them. Our model is deliberately very simple with respect to others available in the literature. Our aim was not to perfectly fit the experimental data but to test the impact of different hypotheses on the channel behavior.

As expected, the transition between closed and open states must be pressure-dependent, favoring the open state when pressure increases. This is in agreement with the large body of literature on mechano-sensitive channels, even if it is now quite well established that the channel is not sensitive to the pressure itself but to its effect on the membrane (e.g., by the curvature induced by pressure) (Cox, Bavi and Martinac, 2019).

The inactivation of the channel is associated with a pressure-independent transition from the open state. This means that inactivation occurs through a different mechanism than the opening-closure (activation/deactivation) of the channel upon pressure application/release and is independent of the membrane condition. Therefore, we propose that it is caused by the blockage of the pore of the open channel by a plug. A possible hypothesis is that ions get trapped inside the channel, needing more time to cross it. A classical example illustrating this phenomenon is the voltage dependent block of NMDA glutamate receptors by Mg^2+^ (Nowak *et al*., 1984). If this is true the inactivation kinetics should change with the transmembane potential (see Figure 3B): the higher the electric field, the easier it should be for ions to cross the membrane. However, we observed voltage independent inactivation of RMA. The chain and ball model proposed for neuronal potassium channels provides an alternative example of voltage independent blockage of the open pore. In this model, a cytosolic domain of the channel acts as a plug. To determine the molecular mechanism of the blockage, a model including structural information about the channel protein is needed (Sukomon, Fan and Nimigean, 2023). Interestingly, the domain structure of DEK1, which has been associated with RMA channel activity (Tran *et al*., 2017b), is compatible with such a chain and ball mechanism. The DEK1 protein includes a large multipass transmembrane domain attached to a cytosolic calpain domain, which could play the role of the “ball” (Kumar, Venkateswaran and Kundu, 2013).

The adaptation of the channel is a poorly understood phenomena, which is in general discarded as a degradation of the channels with time, or a diffusion of channels to the border of the patch. Our results do not agree with such simple hypothesis: the degradation or the diffusion of the channel should be dependent on the time (see Figure 2). We propose here a minimal model to explain the adaptation, with an transition to the adapted closed state exclusively from the open state – at a pressure-independent rate. This implies that adaptation is related to this specific state of the channel, favoring the hypothesis of a biochemical reaction. This is again compatible with the structure of DEK1, as the calpain protease domain is calcium dependent and could be activated upon channel opening and degrade the channel accounting for the irreversible nature of adaptation (Wang *et al*., 2003; Johnson *et al*., 2008; Guerringue, Thomine and Frachisse, 2018; Frachisse, Thomine and Allain, 2020). However, the hypothesis that the calpain mediates activity dependent degradation of RMA cannot be easily tested because deletion of this domain is lethal (Johnson *et al*., 2005; Lid *et al*., 2005).

### The RMA channel behaves like a band-pass filter

An interesting extension of the model is the simulation of RMA responses to oscillating pressure. Indeed, plants are oscillating due to variations of their environments (wind, insect contact, etc.), and have to adapt to these changing environments (Appel and Cocroft, 2014; Veits *et al*., 2019; Tran *et al*., 2021). The simulation of RMA responses to frequency stimulation revealed that the channel behaves as a pass-band filter: the response is higher than the response to static pressure in a broad range of frequencies (around 10Hz to 10kHz). At lower frequency, the response to oscillations is lower than to static pressure. This gain between frequencies <10 Hz and >10 Hz is linked to the inactivation of the channel. At higher frequencies (>1 KHz), the gain associated with frequency stimulation decreases. This effect is linked to the fast close-open transition and has been reported earlier for MSL channels that do not inactivate and exhibit only a low-pass behavior with lower cut-off frequencies (Tran *et al*., 2021). The band-pass filter properties of RMA channels should be integrated with the duration of the stimulations and its adaptation to repetitive stimulations, which was not taken into account in the simulation. Together, these properties may allow RMA to specifically detect oscillations in a specific range of frequencies and duration..

### Integrate the channel in the cytoskeleton cell wall context

In the cellular context, the cell membrane is surrounded by the cell wall while on the inner face the cytoskeleton helps maintain stiffness. Cell wall and cytoskeleton have been shown to be important players in plant mechanotransduction but the role of the membrane squeezed between these two compartments has been for a long time ignored (Colin and Hamant, 2021). Considering the activation inactivation properties of RMA channel we might wonder how this channel function in planta. For a slow stimulation, for instance due to the tension imposed by the turgor pressure counterbalanced by the elastic deformation of the cell wall (estimated at 15%, Kell and Glaser, 1993), the channel would generate no or only weak signals considering that activation will be compensated by inactivation. Upon a fast external mechanical stimulus such as wind or being touched by an animal, RMA would trigger a raise in cytosolic calcium which would rapidly decrease due to its inactivation. For fast and repetitive stimulations as those caused by caterpillar feeding on Arabidopsis (Appel and Cocroft, 2014), or Oenothera plants submitted to acoustic signal of 1 kHz from a speaker, or from the bee buzz (Veits *et al*., 2019), RMA channel would be an excellent candidate to transduce vibration into a primary biological signal. With both sources of stimulation, speaker or bee in the range of kHz, an increase of sugar content in the nectar of Oenothera flowers within 3 minutes is reported, while flowers did not respond to acoustic stimulations at 35 kHz, the sound frequency corresponding to a bat flight (Veits *et al*., 2019). Yet, these studies still do not provide knowledge on the physical mechanism connecting plant vibration with biological response (Son *et al*., 2024). Considering that cell wall is the stiff vibrating component of the plant and the band pass filter property of RMA, this channel is a candidate to consider. The attenuation of the calcium signal due to the adaptation of the channel might be a process to avoid an excess of cytosolic calcium. At the subcellular scale, the generation and sensing of membrane curvature is essential in endo- and exocytosis processes. In addition, this process is accompanied by a local increase in Ca^2+^ concentration. Then RMA could also be a potent actor of this physiological process (Frachisse, Thomine and Allain, 2020; Audemar, Guerringue, Frederick, Melogno, *et al*., 2023; Howell *et al*., 2023).

## Acknowledgements

Y. Guerringue was funded by an allocation from Ecole Normale Superieure de Lyon. This work benefits from the support of the LabEx Saclay Plant Sciences (SPS; ANR-10-LABX-0040-SPS).

## Competing interests

None declared.

## Author contributions

YG and J-MF designed the research. YG carried out the experiments, analyzed the data and prepared the figures. J-MA provided channel modelling. J-MF, J-MA and ST provided the assistance with experiments and contributed with data interpretation. J-MF and J-MA wrote the manuscript with input from YG and ST.

## Supplementary Data

**Figure S1.**
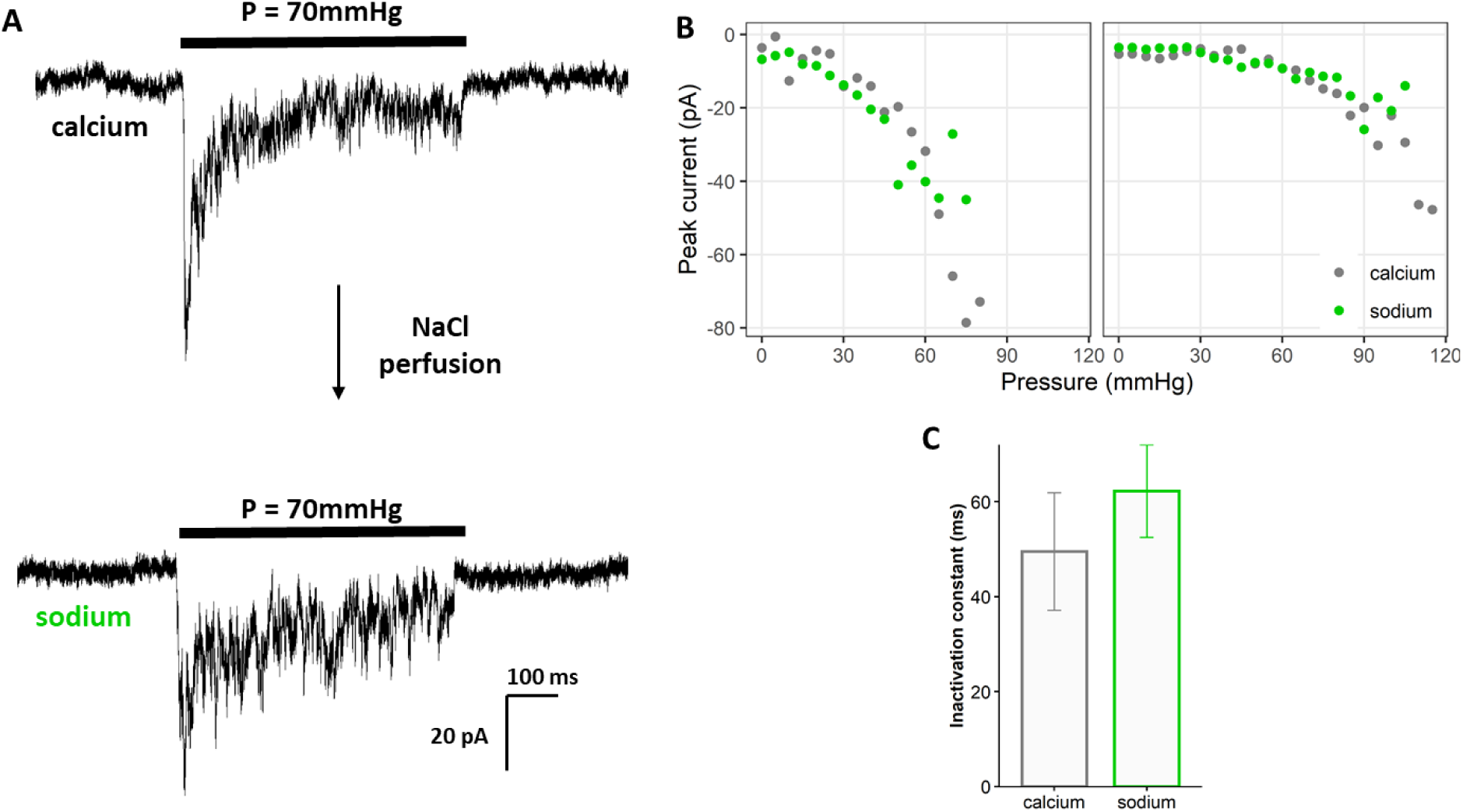
RMA conductance to Na^+^ is similar to Ca^2+^. (A) Recording of the RMA current in the same patch under a 70 mmHg-pressure pulse in standard 50 mM CaCl_2_condition (up) and after the perfusion of a bath solution containing 100 mM NaCl . (B) Peak current at different values of pressure for two independent patches before (*grey*) and after (*green*) the perfusion of the Na^+^ solution. No difference in peak intensity has been detected between the two conditions, suggesting that the conductance to Na^+^ and Ca^2+^ are similar. (C) Average inactivation constant measured at different pressure values (Mean ± SE, n = 5 patches, Wilcoxon test p-value = 0.81).

**Figure S2.**
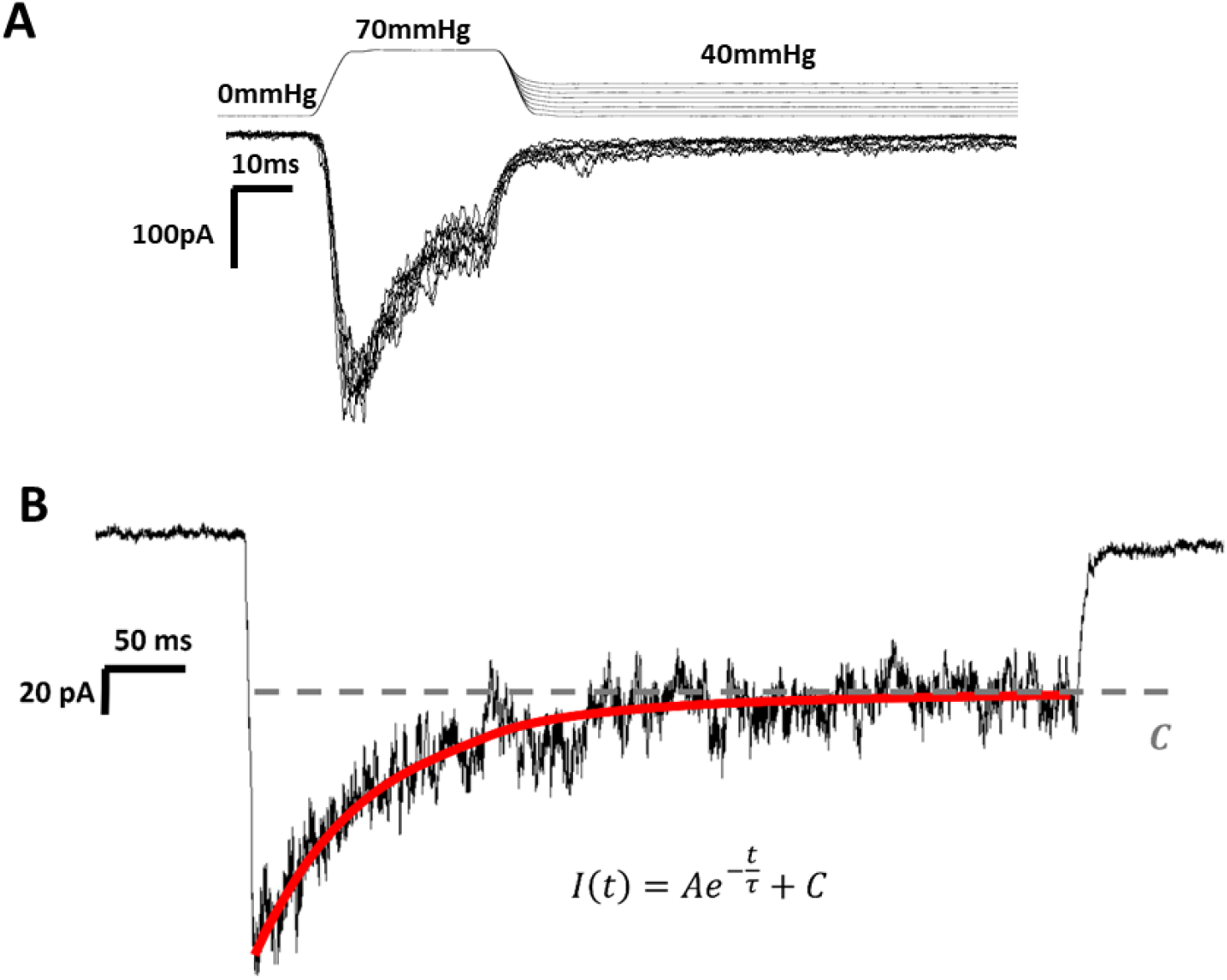
Dynamics of RMA. (A) Pressure protocol to test pressure-dependence of the deactivation consisting of a double pressure pulse composed of one pulse at the same pressure and a second at decrease pressure from 40 to 0 mmHg. We could not detect a difference in deactivation kinetics with different pressure values. As in the case activation, the rate of deactivation is faster or similar to the time response of the pressure clamp system. (B) Exponential fit (*red*) of the time-dependent decrease of RMA current during a pressure pulse at 100 mmHg (black). The equation of the fit is shown below. Ae is the amplitude of the inactivating current, C is the amplitude of the residual current and *τ* is the inactivation constant.

**Figure S3.**
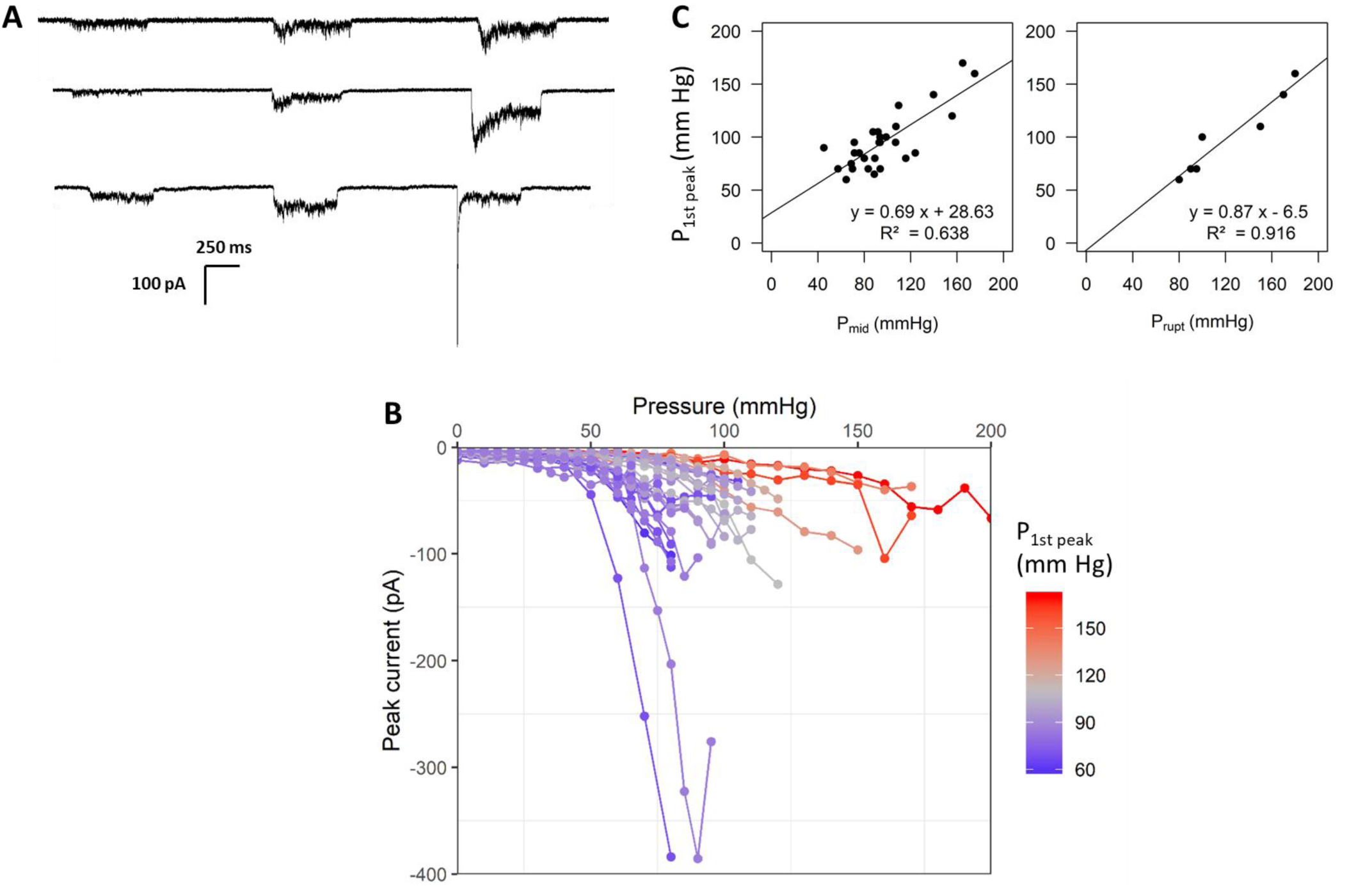
The pressure at the first peak (P_1st peak_) correlates with RMA mid activation pressure (Pmid) and the membrane rupture pressure (Prupt). (A) Recordings of the RMA conductance in three independent patches under square pulses of increasing pressure before and at the first peak. (B) Peak current at different pressure values. Each curve corresponds to one independent patch and is colored according to the pressure at first peak *P_peak_* (in mmHg, n = 27?)). (C) The pressure at first peak *P_1st peak_* was correlated to the pressure at mid-activation *P_mid_* (n= 27) and the rupture pressure *P_rupt_*.(n =7). Linear correlation fit was represented by a straight line which equation is indicated with the correlation coefficient.

**Figure S4.**
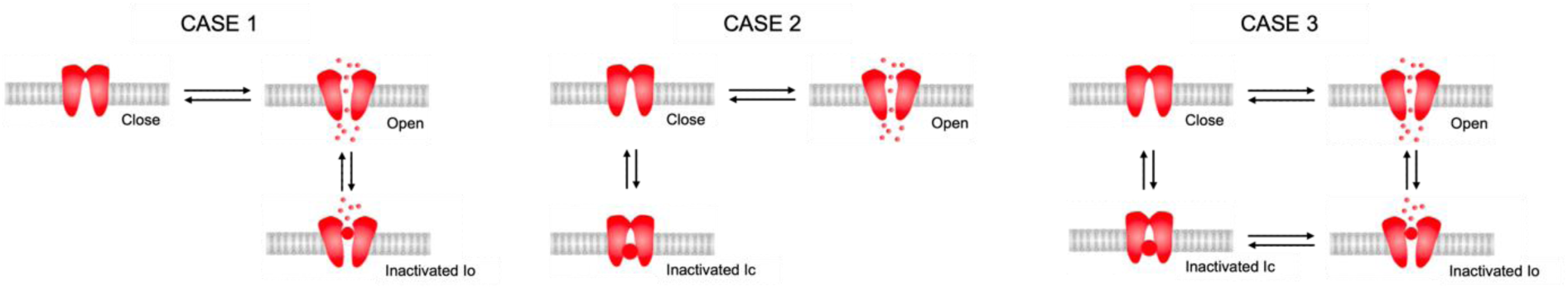
Sketch of the 3 differents models for the inactivation. The first case corresponds to an inactivation which is connected to the open state. The second case corresponds to an inactivation connected to the close state addressing the hypothesis of the adaptation due to the inactivation. Case 3 combines the two previous approaches.

**Supplementary Data S5 – Equations and constants of the model**

The channel is described by 4 states (see Figure 5A): closed (C), open (O), inactivated (I), and adapted (A).

We consider a total number of channels embedded in the membrane *N*_*tot*_. The number of channels in a given state (*⍺*) is *N*_*⍺*_. The fraction of channels in the state (*⍺*) is

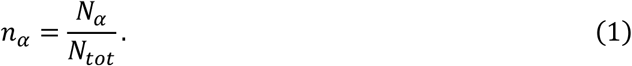

The sum of the channel fractions in the different states is therefore equal to 1:

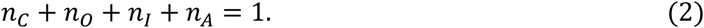

The evolution of the channel fractions is given by the equations:

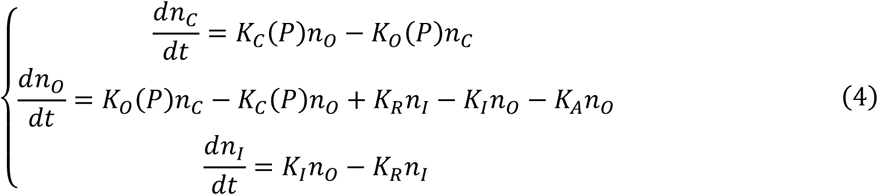

The constraints on the sum of the channel fractions (Eq. 2) makes useless the equation for the evolution of the adapted channel fraction:

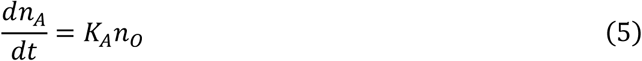

In our model, only the opening and closing rates *K*_*O*_ and *K*_*C*_ change with the tension on the channel Σ. Experimentally, we do not access to this tension. So, we assumed that the tension is proportional to the applied pressure Σ = αP.

For the evolution of the rates with the tension, we assumed the classical exponential behavior (Tran *et al*., 2021):

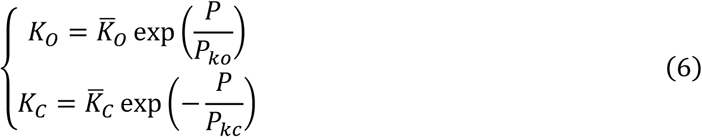

The constants *P*_*ko*_ and *P*_*kc*_ have to be determined for each experimental conditions, as they will depend on the proportionality coefficient between the tension and the pressure.

The experimental observation is the current versus time, which is proportional to the total number of open channels:

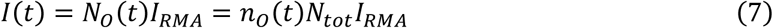

where *I*_*RMA*_ is the current generated by the opening of a single channel. Comparison with the experiments allows us to determine the quantity *N*_*tot*_ *I*_*RMA*_.

The experimental protocol consists of periods under pressure, preceded (and followed) by pressure-free periods. In each period, the pressure is kept constant. For simplicity, we assumed that the pressure rose and decreased instantaneously: this enabled us to assume piece-wise functions for the pressure, and thus for the kinetic constants.

During a period on which the pressure doesn’t change, the constants do not change also. Therefore, if we know the initial channel fractions, we can find an analytical solution for the fraction evolution with time (Eq. 4) during the full period.

Our procedure to reproduce the experimental data is then to first obtain the experimental times at which the channel is under pressure, and at which pressure, and the times at which is not under pressure. This defines a series of periods, of known pressure and duration.

To initialize the different fractions, we used the stationary solution of Eq. 4, under the assumption that there is no adaptation (so *K*_*A*_ = 0).

Then, starting from these initial fractions, we used the analytical solution of Eq. 4, including the adaptation, to determine the evolution of the fraction with time during the first period. The final values of the fractions are used as initial conditions for the next period. And we keep going up to the end of the experiment.

We then compare the intensity versus time for increasing pressure steps (see Fig. 5B-D). In the same experiment, a series of periods at increasing pressures is performed (see Fig. 5B-C), and a series of periods at the same pressure (see Fig. 5D).

The different parameters of the model are then manually optimized to reproduce the experimental data. We didn’t perform a sensitivity analysis, as our aim is not to determine the different coefficients, but to determine the simplest model which mimics the data.

The Supplementary Table 1 shows the parameters obtained for the experiment reproduced in Figure 5.

**Supp. Table 1:** values of the parameters used to reproduce the experiment in Figure 5.

| $\bar{K}_O$ | $P_{ko}$<br>(mmHg) | $\bar{K}_C$ | $P_{ko}$<br>(mmHg) | $K_I$ | $K_R$ | $K_A$ | $N_{tot}I_{RMA}$<br>(pA) |
| --- | --- | --- | --- | --- | --- | --- | --- |
| 0.01 | 35 | 750 | 42 | 0.15 | 0.003 | 0.0055 | 550 |

Using these parameters, one can estimate the initial number of channels around 275. The inactivation time of the channel is around 7ms, of the same order of magnitude than the experimental observations (around 20ms). The adaptation time is around 180ms, so typically 10 times larger.

The values of the characteristic pressure of the variation of open and closing rates do not carry any real meaning without the knowledge of the proportionality constant with the tension on the channel.

## Notes

### Competing Interest Statement

The authors have declared no competing interest.

